# A four-point molecular handover during Okazaki maturation

**DOI:** 10.1101/2022.08.02.502465

**Authors:** Margherita M. Botto, Alessandro Borsellini, Meindert H. Lamers

## Abstract

DNA replication introduces thousands of RNA primers into the lagging strand that need to be removed for replication to be completed. In *Escherichia coli* when the replicative DNA polymerase Pol IIIα terminates at a previously synthesized RNA primer, DNA Pol I takes over and continues DNA synthesis while displacing the downstream RNA primer. The displaced primer is subsequently excised by an endonuclease, followed by the sealing of the nick by a DNA ligase. Yet how the sequential actions of Pol IIIα, Pol I, endonuclease and ligase are coordinated is poorly defined. Here we show that each enzymatic activity prepares the DNA substrate for the next activity, creating an efficient four-point molecular handover. The cryo-EM structure of Pol I bound to a DNA substrate with both an upstream and downstream primer reveals how it displaces the primer in a manner analogous to the monomeric helicases. Moreover, we find that in addition to its flap-directed nuclease activity, the endonuclease domain of Pol I also specifically cuts at the RNA/DNA junction, thus marking the end of the RNA primer and creating an 5’ end that is a suitable substrate for the ligase activity of LigA once all RNA has been removed.

## INTRODUCTION

In *E. coli*, DNA replication is performed by the DNA polymerase III holoenzyme, a large multi-protein complex that synthesizes DNA at speeds of up to 1000 nucleotides (nt) per second^1,2^ and synthesis lengths of up to 100,000 base pairs (bp) per binding event^3^. Due to the opposite polarity of the two DNA strands, DNA replication occurs asymmetrically, where the leading strand is synthesized in a continuous fashion while the lagging strand is synthesized in short fragments of 500-1000 bp termed Okazaki fragments^4,5^. Hence during replication of the 4.6 million bp of the *E. coli* genome^6^, the lagging strand is created in up to 9000 fragments. As the replicative DNA polymerase, the α subunit of the DNA polymerase III holoenzyme (Pol IIIα) cannot start DNA synthesis *de novo*, each Okazaki fragments is initiated by a 10-15 nt RNA primer that is synthesized by the DNA primase DnaG^7,8^. To complete DNA replication, these thousands of RNA primers are removed from the DNA by the consorted action of Pol IIIα, DNA polymerase I (Pol I), the endonuclease domain of Pol I, and a DNA ligase that together remove the RNA primer, resynthesize the gap and seal the nick between two adjacent Okazaki fragments.

The roles of the individual proteins and subunits has been well described (for a review, see^9^). The bulk of the DNA is synthesized by Pol IIIα that when bound to the processivity clamp β is a fast and highly processive enzyme^10,11^. Yet, Pol IIIα abruptly comes to a halt when it encounters a downstream RNA primer of the previously synthesized Okazaki fragment and disengages from the DNA^12^. DNA synthesis is then continued by Pol I that is capable of strand displacement of the downstream RNA primer, while simultaneously extending the upstream DNA strand^13,14^. The displaced RNA primer is then removed by the N-terminal domain of Pol I that is a flap-directed 5’-3’ exo/endonuclease^15–18^. Once the RNA primer has been removed, a DNA ligase seals the nick between the two adjacent DNA fragments^19,20^.

While the basic roles of the different proteins and subunits are known, several fundamental questions remain about how the removal of the RNA primer is orchestrated: i) how is Pol I able to continue DNA synthesis where Pol IIIα comes to a halt? ii) how does Pol I detect the end of the RNA primer and not continue strand displacement into the DNA section of the Okazaki fragment? iii) how is the endonuclease activated to cut the displaced RNA flap and directed to remove all of the RNA primer? iv) what prevents the ligase from ligating the DNA to the RNA primer? and finally, v) how are the sequential activities of Pol IIIα, Pol I, endonuclease, and ligase organized in time and prevented from counteracting each other’s activities?

Here we present the first structure of Pol I engaged with both an upstream (the extended) primer and a downstream (the displaced) primer. The cryo-EM structure of Pol I engaged with the both primers reveals how Pol I acts in a manner analogous to monomeric helicases such as UvrD and RecQ^21,22^, where the fingers domain of Pol I acts similar to the strand separating pin of the helicases, while the primer extension of Pol I acts as a motor that drives the translocation of the template strand, analogous to the ATPase domains of the monomeric helicases. The DNA itself is kinked by 120° positioning the ingoing and outgoing DNA sections at an almost perpendicular angle. We furthermore present the biochemical analysis of the sequential activities of the four enzymes involved in the Okazaki maturation. We show that the endonuclease domain of Pol I is a unique RNA-DNA directed endonuclease that specifically introduces a nick at the RNA-DNA junction, thus marking the end of the RNA primer. In addition, we show that LigA, the dominant *E. coli* DNA ligase, cannot ligate RNA to DNA, but will act as soon as a DNA-DNA junction is available. The ligation of the two Okazaki fragments results in the eviction of Pol I from the DNA, which has a 20-fold reduced affinity for continuous dsDNA compared to nicked DNA, thus completing the Okazaki fragment maturation.

Hence, our work reveals that the activities of the different proteins act as a ‘molecular relay race’, in which each activity prepares the DNA/RNA substrate for the next enzymatic activity to take over. This way, the removal of the thousands of RNA primers from the lagging strand introduced during DNA replication is an efficient process that ensures that an intact DNA is delivered for the next generation of cells.

## RESULTS

### Molecular handover between Pol IIIα and Pol I

When the replicative DNA polymerase Pol IIIα encounters a previously synthesized Okazaki fragment it terminates DNA synthesis^12^, which is then continued by Pol I^13,14^. To determine how Pol IIIα and Pol I trade place on the DNA we measured their polymerase activity on a DNA substrate containing an upstream primer (the extended primer) in the absence or presence of an RNA or DNA downstream primer (the displaced primer), separated by a 15-nucleotide gap. For the reaction, the β-clamp was added to both polymerases, and Pol IIIα was supplemented with its 3’-5’ proofreading exonuclease ε (creating Pol IIIαε), while Pol I contains its 3’-5’ proofreading exonuclease domain within the same polypeptide. In the absence of a displaced primer, both polymerases synthesize to the end of the DNA template (Fig. 1a). Similarly, in the presence of a displaced primer DNA synthesis by Pol I is unaffected and continues until the end of the template strand (Fig. 1b), in agreement with its described strand displacement activity^13,14,23^. In contrast, Pol IIIαε comes to a halt at the start of the displaced primer, irrespective of the RNA or DNA nature of the primer as was previously shown^12^. Next, to determine if Pol IIIαε leaves a gap between the extended primer and the displaced primer, we used a displaced DNA primer with a 5’ phosphorylated end that is required for DNA ligase activity and added the *E. coli* DNA ligase LigA after initiation of DNA synthesis (Fig. 1c). In the presence of Pol IIIµε and LigA, the extended primer is ligated to the displaced primer to create a fully extended product of 69 nt, indicating that Pol IIIµε continues DNA synthesis up to the very last nucleotide before handing over to Pol I. However, during DNA replication, ligation of the newly synthesized DNA segment and the downstream RNA primer is prevented as the 5’ end of the RNA primer starts with a di- or tri-phosphate nucleotide^7^ that is not a suitable substrate for ligase^24^. Instead, the RNA primer is displaced by the strand displacement synthesis of the Pol I.

**Fig. 1.**
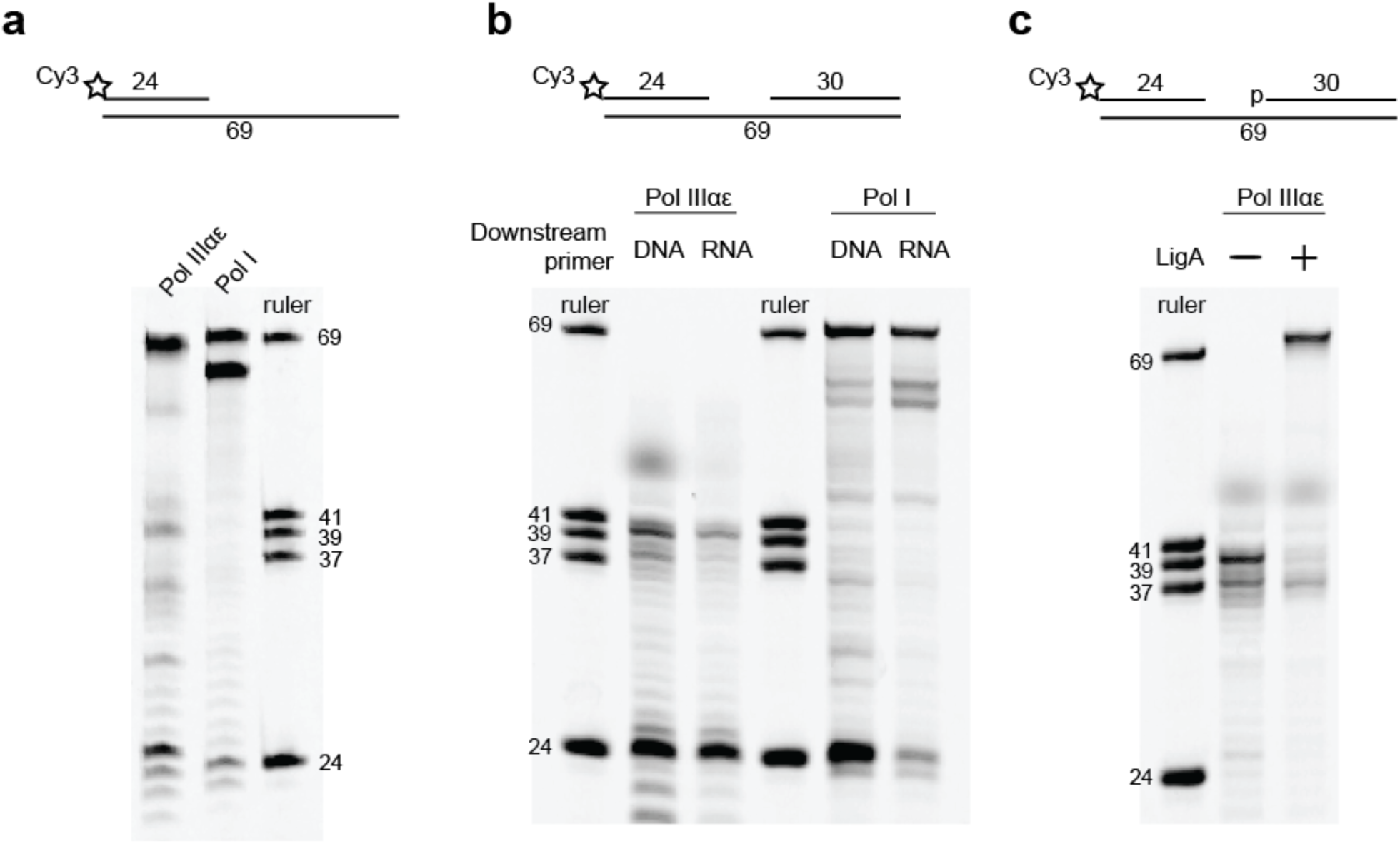
Comparison of Pol IIIαε and Pol I primer extension activity. **a**, DNA synthesis of Pol IIIαε and Pol I on a 69 nt template strand with a single upstream primer. Right lane (marked ‘ruler’) shows DNA substrates of different lengths as indicated. **b**, DNA synthesis of Pol IIIαε and Pol I on the same substrate as panel A complemented with a DNA or RNA downstream primer. **c**, DNA ligase activity by LigA after primer extension between the upstream and downstream primer by Pol IIIαε. The downstream primer contains a 5’ phosphate group needed for ligase activity. All the reactions were performed in presence of the β clamp.

### Cryo-EM structure of Pol I bound to an Okazaki fragment

To understand how Pol I is able to continue DNA synthesis in the presence of a displaced primer, we determined the cryo-EM structure of Pol I bound to a DNA substrate containing both an upstream and downstream primer with a six-nucleotide single stranded flap to a resolution of 4.3 Å (Fig. 2 and Extended Data Fig. 1). Although full length Pol I was used (Fig. 2c), no density was observed for the endonuclease domain, indicating that it is flexible in this structure. Therefore, in an attempt to increase the resolution of the cryo-EM map, we used a version of Pol I that lacks the N-terminal endonuclease domain (also known as Klenow fragment, here indicated as Pol I^KL^) to collect a large cryo-EM data set of over 11,000 images and over 500,000 initial particles. This dataset yielded two structures of Pol I bound to the DNA substrate, to a final resolution of 4.0 and 4.1 Å. Hence, the use of Pol I^KL^ did not result in an improved resolution of the map, but the increase in the number of particles did allow for the separation of the two different conformations that Pol I takes on under these conditions. The first structure shows Pol I^KL^ with the 3’ terminal base pair of the extended primer in the polymerase active site and the 5’ terminal base pair of the displaced primer stacked against the fingers domain. The second structure shows the DNA translated by one nucleotide in the direction of DNA synthesis, pushing the extended primer further out and separating one additional nucleotide from the displaced primer. Furthermore, the translation of the DNA creates an unpaired nucleotide in the polymerase active site as no dNTPs were added to the sample. The one nucleotide translation of the DNA is accompanied by a canonical movement of the fingers domain^25–27^ that creates a more open active site for an incoming nucleotide. In contrast, the non-translated structure shows a closed active site with the the O-helix of the fingers domain stacked against the 3’ terminal base pair of the extended primer.

**Fig. 2.**
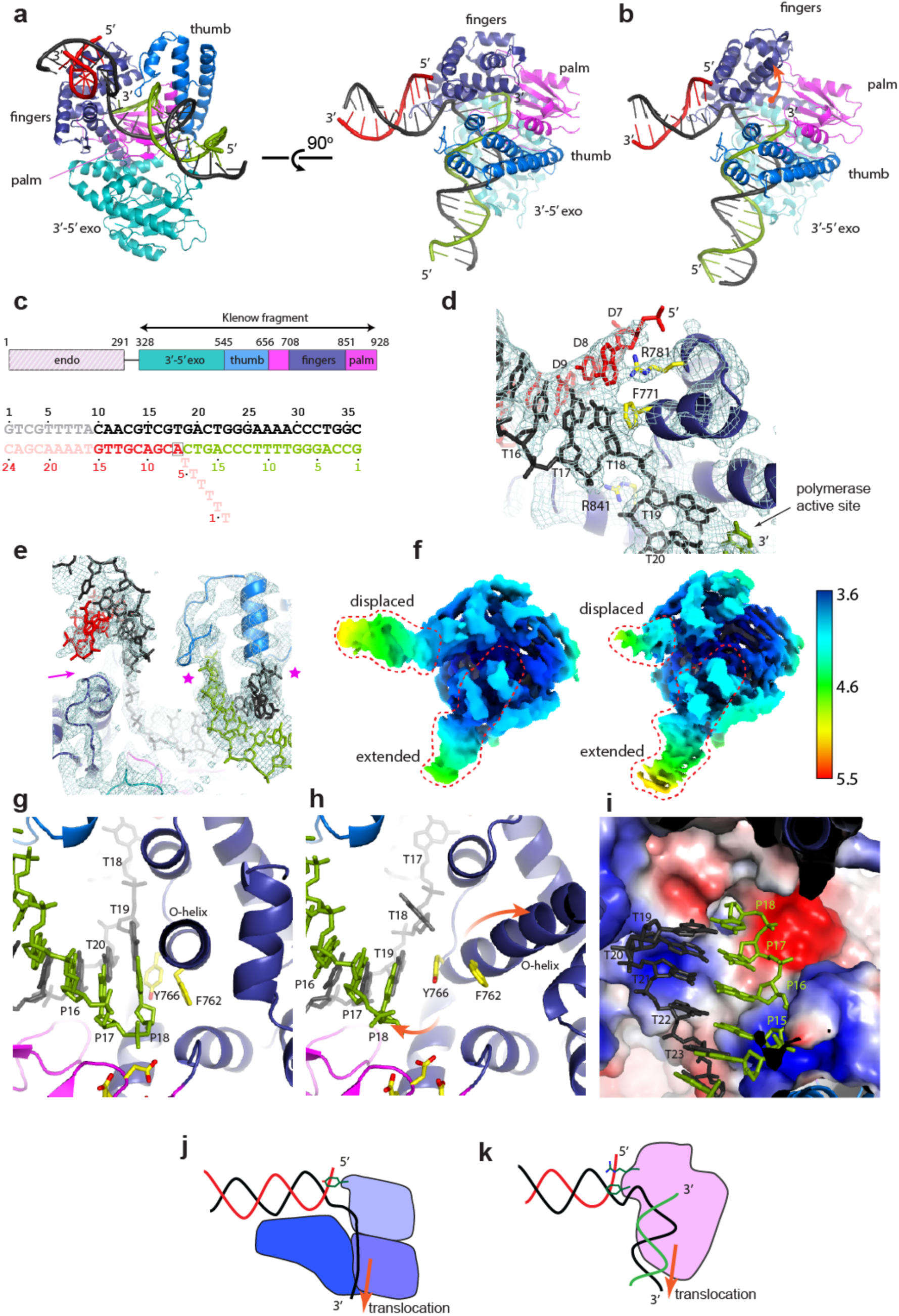
Cryo-EM structures of Pol I bound to an upstream and downstream DNA substrate. **a**, Front view (left) and top view (right) of closed Pol I structure with different domains marked. Continuous, template DNA strand in black color, extended primer strand in green color, and displaced primer strand in red color. **b**, Top view of the open Pol I structure. Movement of the fingers domain indicated with the orange arrow. **c**, Schematic view of Pol I protein and DNA substrate used for cryo-EM structure determination. Different domains are indicated. Striped coloring of the endonuclease domain indicates the absence of the domain in the structure. Full DNA substrate with template strand in black, extended primer in green, and displaced primer with single stranded flap in red color. Sections of the DNA not observed in the cryo-EM map are marked in transparent coloring. **d**, Close up of the interaction of the fingers domain and the displaced section of the DNA. Cryo-EM map is shown in light blue mesh. Three key residues that interaction with the displaced DNA are shown in yellow sticks. Labeling of the nucleotides in the template strand (T) and displaced strand (D) follows the numbering of panel c. **e**, Close up of the protein-DNA interactions of the displaced and extended DNA section. Cryo-EM maps shown in light blue mesh. Pink arrow indicates the limited contact of the displaced section of the DNA with the polymerase fingers domain. Pink stars mark the contacts between the polymerase thumb domain and the extended section of DNA. **f**, Local resolution maps of the closed (left) and open (right) Pol I structures colored from high resolution (blue) to low resolution (yellow/red). Red dashes lines mark the position of the displaced and extended section of the DNA. **g**, Close-up of the polymerase active site in the closed Pol I structure. Labeling of the nucleotides follows the same numbering as in panel c. Two aromatic residues that interact with the terminal base pair and the three glutamates/aspartates of the catalytic triad are shown in yellow sticks. **h**, Same view for the open Pol I structure. Movement of the DNA and the O-helix is marked with the two orange arrows. **i**, Electrostatic potential plot of the polymerase active site of the closed structure. A strong negatively charged patch is located on the terminal nucleotides of the extended primer strand. Parts of thumb and fingers domain were omitted for clarity. **j**, Schematic drawing of the mechanism of strand displacement by monomeric helicases such as UvrD. **k**, Schematic drawing of the mechanism of strand displacement by Pol I.

Overall, the closed (non-translated) and open (translated) structures are very similar and both show a striking ∼120° kink of the DNA that places the 3’ end of the extended primer in the polymerase active site while the 5’ end of the displaced primer interacts with the fingers domain. While multiple structures of Pol I bound to an extended primer exist^28–31^, this is the first time that also the displaced primer is visualized. The position of the extended primer is identical to that of the previous structures of Pol I bound to a DNA substrate, while the position of the displaced primer is in agreement with a recent study where Förster resonance energy transfer (FRET) was used to predict its location^32^. Importantly, our structure reveals for the first time the molecular details of the interactions between Pol I and the displaced primer (Fig. 2d), which are best defined in the closed structure. The first base pair of the displaced section interacts with F771 and R781 that stack on the template and primer base, respectively. R841, positioned under the template strand, acts as a pivot point over which the DNA is bent by 120 °. F771 and R841 were previously shown to be important for the strand displacement activity of Pol I^14^. Remarkably, the displaced section of the DNA has only limited contact with the protein, in contrast the extended section of the DNA that has extensive contacts (Fig. 2e). Concordantly, the cryo-EM map for the displaced section of the DNA is poorly resolved and only nine of the 18 base pairs can be placed (Fig. 2a and 2f - left panel). In contrast, the cryo-EM map for the extended DNA section is well resolved in which all 18 base pairs can be placed. Furthermore, upon translocation of the DNA in the open structure, the displaced DNA section shows a weaker cryo-EM map, indicating even less interaction between protein and the displaced DNA in this conformation (Fig. 2b and 2f right panel, Supplemental Video 1).

In the polymerase active site of the closed structure, the 3’ terminal base pair is stacked against the O-helix (Fig. 2g), while in the open structure the DNA and O-helix have moved in opposite directions (Fig. 2h). During this movement, two aromatic residues, phenylalanine 762 and tyrosine 766 trade places with respect to their interaction with the DNA. In the closed structure, phenylalanine 762 stacks onto the sugar ring of the 3’ terminal nucleotide of the extended strand, while tyrosine 766 is tucked away under the DNA (Fig. 2g). In the open structure, phenylalanine 762 is separated from the 3’ terminal nucleotide by ∼6 Å, while tyrosine 766 now stacks on the base of the template strand (Fig. 2h). Due to the one-nucleotide translation of the template strand, a single unpaired nucleotide is created in the polymerase active site (nucleotide 18 of the template strand) as no dNTPs were added to the sample. In the closed structure, this unpaired thymine was paired with an adenosine nucleotide in the displaced strand, that is now become part of the single stranded flap and no longer visible in the cryo-EM map.

The translation of the DNA, the movement of the O-helix, and unpairing of the first base pair of the displaced DNA section raise the question how the translation is achieved in the absence of dNTPs. Inspection of the polymerase active site reveals a large negatively charged patch at the position of the 3’ terminus of the extended primer strand that could act as a repulsive force to the negatively charged backbone of the DNA (Fig. 2i). The translation of the DNA by one nucleotide moves the 3’ end of the primer strand away from the negative patch that could therefore contribute to the translation of the DNA.

A molecular morph between the two structures gives further insight into how the DNA moves through the protein during DNA synthesis (Supplemental Video 2). As the extended strand moves out of the active site, it pulls along the continuous template strand. This in turn, pulls the displaced strand into the fingers domain, resulting in the displacement of the downstream primer and the lengthening of the single stranded flap. The translocation of the template strand pulls the next template base into the polymerase active site where it is ready to pair with the next incoming nucleotide. The opening of the O-helix further facilitates the access of the new nucleotide into the polymerase active site.

Remarkably, the mechanism of strand separation in Pol I can be compared to that of the monomeric helicases such as RecQ and UvrD (reviewed in ^21,22^). Here, the two ATPase domains create the driving force that pulls the continuous DNA strand through the protein, while a ‘strand separation pin’ splits the complementary strand from the template strand (Fig. 2j). In an analogous manner, in Pol I, DNA synthesis in the polymerase active site acts as the driving force for the translocation of the template strand, while the fingers domain acts as the strand separation pin that splits the downstream primer from the template strand (Fig. 2k).

### Pol I endonuclease and polymerase domain compete for DNA binding

As Pol I performs strand displacement DNA synthesis it generates a single stranded flap that becomes a substrate for the endonuclease domain. Yet how the endonuclease domain gains access to the single stranded flap is not known. The DNA substrate used for the cryo-EM structures contains a six-nucleotide single-stranded in the displaced primer but these are not observed in the cryo-EM map, indicating that the single stranded flap is flexible. However, based on the position of the 5’ of the displaced primer from which the single stranded flap would continue, it would stand out on top of the fingers domain. Here it could be reached by the endonuclease domain that is connected to polymerase via a ∼35 amino acid unstructured linker (Fig. 3a). However, as mentioned above, in our structure of full length Pol I no additional density was observed that can be attributed to the endonuclease domain, indicating that is flexible in this structure. The predicted flexibility of the endonuclease domain is in agreement with the crystal structures of *Thermus aquaticus* Pol I^33^ and that of *Mycobacterium tuberculosis* Pol I^31^ in which the endonuclease domain is found in two different positions (Extended Data Figs. 2b-c).

**Fig. 3.**
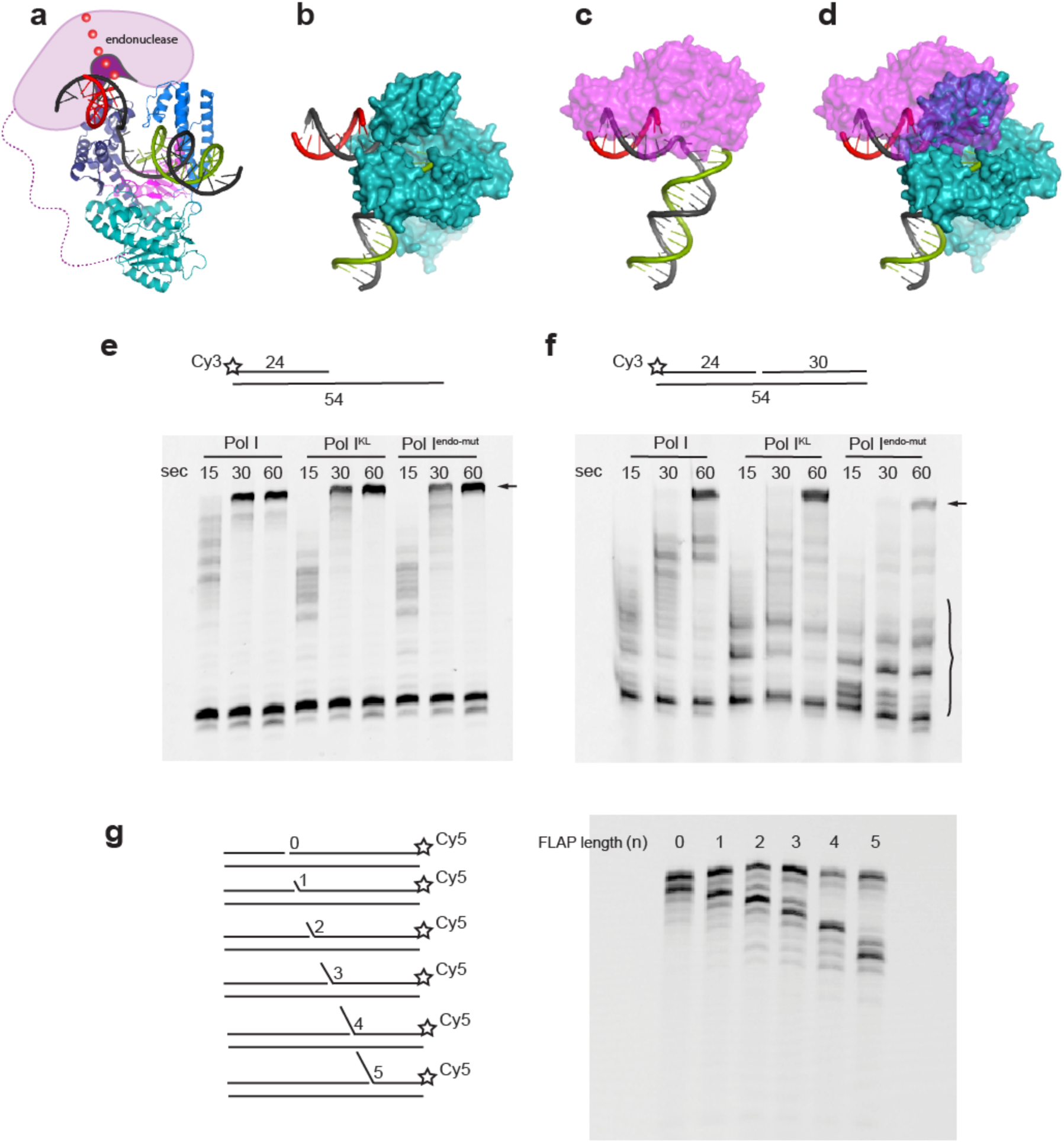
Alternating activities of endonuclease and polymerase domains. **a**, Front view of the Pol I-DNA structure with the predicted position of endonuclease domain marked in pink/purple cartoon. The approximate position of the single stranded flap is indicated by the red spheres. **b**, Cryo-EM structure of Pol I bound to an Okazaki fragment. Pol I is shown in blue surface, the template strand in black, the extended primer in green, and displaced primer in red. View similar to Fig. 2a-b. **c**, Model of Pol I endonuclease domain bound to the displaced primer. Endonuclease domain shown in transparent magenta surface. For modelling details see Extended Data Fig. 2. **d**, Combined structures of Pol I and endonuclease domain reveal a large overlap between the two structures. **e**, Primer extension activity on a primed substrate by Pol I, Pol I Klenow fragment (Pol I^KL^), and Pol I with inactive endonuclease domain (Pol I^endo-mut^). Arrow marks full length product. **f**, Primer extension activity on a double primed substrate. Arrow marks full length product and bracket marks incomplete products. **g**, Endonuclease activity of Pol I on DNA substrates with different lengths of single stranded flaps (0-5 nt).

To gain insight into how the endonuclease domain might engage with the single stranded flap, we modeled how the endonuclease domain could interact with the single stranded flap that emanates from the top of the fingers domain. First, a model of *E. coli* Pol I endonuclease domain was generated using AlphaFold^34^ that overlays well with the endonuclease domains of *T. aquaticus* and *M. smegmatis* Pol I^31,33^, (Extended Data Fig. 2d). Next, as currently no structure has been determined of a Pol I endonuclease domain bound to DNA, we used crystal structures of the structurally related flap-endonucleases FEN1 and T5 endonuclease bound to a DNA substrate^35,36^ to model the binding of the Pol I endonuclease domain to the downstream DNA substrate (Extended Data Figs. 2e-f). Finally, we then used the DNA substrate of the FEN1 and T5 endonuclease structures as a guide to position *E. coli* Pol I endonuclease domain onto the displaced DNA section of our Pol I cryo-EM structure (Fig. 3b-c). The resulting superimposition of the endonuclease onto the downstream section of the DNA in our Pol I structure results in a major clash with the fingers domain of the polymerase (Fig. 3d). This suggests that in order for the single stranded flap to be cut by the endonuclease domain, the polymerase domain may temporarily dissociate from the DNA. Indeed, previous biochemical and single molecule studies have shown that the endonuclease and polymerase domain alternate on the DNA^37,38^. The predicted clash of the two domains furthermore explains the absence of the endonuclease domain from our structure as it cannot co-exist with the polymerase on the same DNA.

As the two domains compete for the same substrate, we wondered if the action of the endonuclease domain can affect DNA synthesis of the polymerase domain. For this we compared three versions of Pol I: full length Pol I, Pol I with the endonuclease domain deleted (Klenow fragment: Pol I^KL^) and full length Pol I in which two active site residues of the endonuclease domain (D115 and D140) were mutated to alanine to render it inactive (Pol I^endo-mut^). On a DNA substrate without a downstream primer, all three proteins synthesize DNA in a similar manner, indicating that all version retain full polymerase activity (Fig. 3e).

In the presence of a downstream primer, both Pol I and Pol^KL^ show a somewhat reduced activity, indicating that the downstream primer slows down polymerase. Importantly, Pol I^endo-mut^ shows an even more pronounced slowing down of polymerase activity with fewer full-length products and more intermediate fragments of the extended primer (Fig. 3f). This suggests that the mutated endonuclease binds to the displaced strand, but as it is not able to perform the incision, it remains bound to the displaced strand and thus slows down the polymerase. This therefore supports the notion that the polymerase and endonuclease domain alternate on the DNA. However, the wild-type protein appears unaffected by the presence of the endonuclease domain indicating that the alternating activities do not slow down the polymerase.

Finally, to determine if the endonuclease requires a specific length of the single stranded flap before it can act, we measured the endonuclease activity on a series of DNA substrates that mimic the progression of DNA synthesis by increasing the length of the extended primer and the length of the single stranded flap on the downstream primer (Fig. 3g). Nucleotides were omitted in order to isolate the endonuclease activity from polymerase activity. On all the substrates similar activity is observed, indicating that the endonuclease domain does not have a specific length requirement. Moreover, on all substrates the cut occurs one nucleotide further than the end of the single stranded flap, in agreement with earlier reports^17,18^. This is also consistent with our cryo-EM structures that show that even in the absence of dNTPs Pol I is able to translate the DNA by one nucleotide, thus enabling the endonuclease to cut one more nucleotide into the displaced strand.

### Pol I endonuclease domain is an RNA/DNA-junction specific endonuclease

As all experiments above were conducted with a DNA downstream primer, we also compared the endonuclease incision on an RNA/DNA-hybrid downstream primer that is similar to the natural substrate encountered during Okazaki maturation (Fig. 4a). In the absence of dNTPs, when the polymerase cannot proceed, we find that the endonuclease removes the first one or two nucleotides on both the DNA and RNA/DNA substrate (Fig. 4a). Surprisingly, we also find that on the RNA/DNA substrate, but not the DNA-only substrate, an additional incision takes place at the junction between the RNA and DNA. To our knowledge, this is the first time an RNA/DNA-junction specific endonuclease activity is observed. This novel RNA/DNA-junction directed endonuclease activity is not observed for the isolated endonuclease domain, nor when polymerase and endonuclease domains are added as separate proteins (Fig. 4b) suggesting that polymerase domain and endonuclease work together. However, the RNA/DNA junction specific incision is also observed on substrates with an increasing gap between the extended primer and the downstream primer where the polymerase is positioned further away from the downstream primer (Fig. 4c). This suggests that the requirement of the polymerase domain is mainly to bring the endonuclease to the RNA/DNA substrate, rather than directly affect the downstream RNA/DNA primer. The incision at the RNA/DNA junction is intriguing as it may make the removal of the RNA primer more efficient. By marking the end of the RNA primer it could help prevent the continuation of the strand displacement into the DNA section of the downstream primer.

**Fig. 4.**
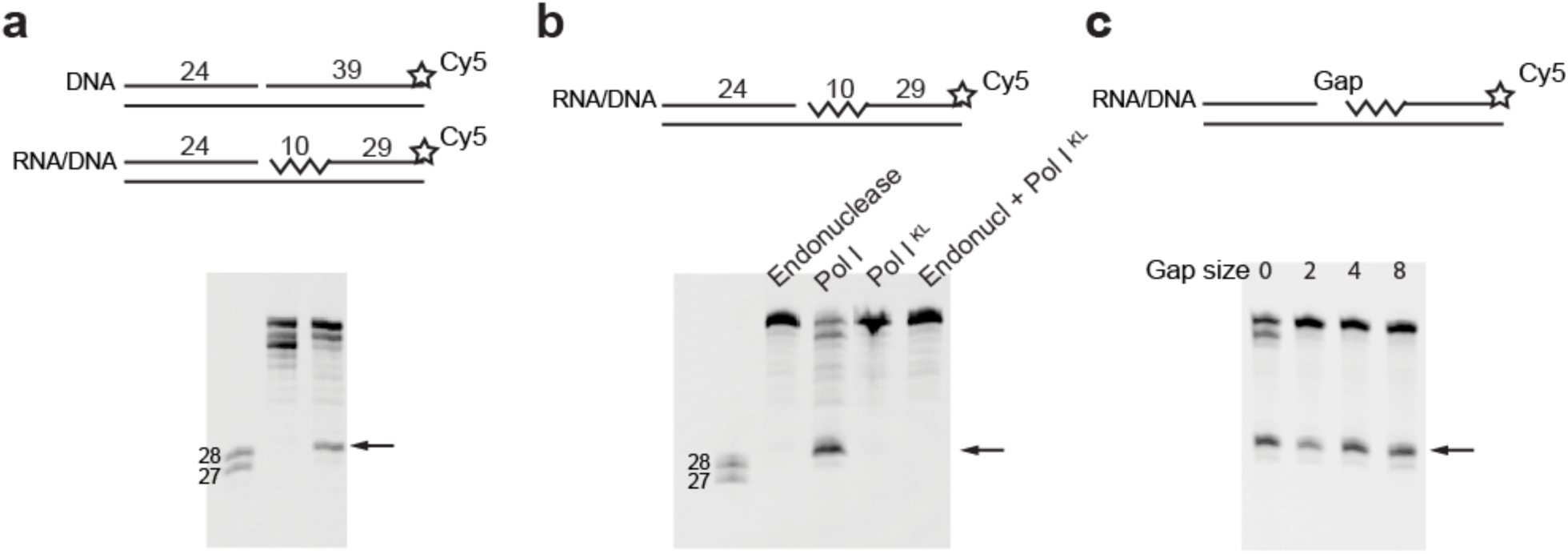
Pol I endonuclease cuts at the RNA/DNA junction. **a**, Endonuclease activity of Pol I on an all DNA downstream primer (‘DNA’) and a RNA/DNA downstream primer (‘RNA/DNA’) containing ten RNA nucleotides (indicated by the zig-zag line) and 29 DNA nucleotides. Arrow marks the 29 nt product after incision at the RNA/DNA junction. **b**, Endonuclease activity on the RNA/DNA substrate of the isolated endonuclease domain, full length Pol I, the isolated Pol I^KL^ domain and the isolated endonuclease domain in combination with the Pol I^KL^ domain. **c**, Endonuclease activity on RNA/DNA substrate with an increasing gap size between the upstream primer and downstream primer.

### LigA seals the nick when the RNA is completely removed

As shown above, when Pol IIIα terminates DNA synthesis at the downstream primer it leaves a nick that can be ligated when the downstream primer is DNA (Fig. 1c). However, during DNA replication, each DNA segment is preceded by an RNA primer that contains a triphosphate tail at its 5’ end^7^ that is not a compatible substrate for ligase activity^24^. However, the flap directed action of the endonuclease does create clean 5’ ends that could potentially become a target for a ligase.

Therefore, to determine if the ligation can also occur to a downstream RNA primer, we tested the two *E. coli* ligases, LigA and LigB, on a series of downstream primers with different number of RNA nucleotides (Fig. 5a). On a DNA:DNA substrate, LigA efficiently ligates the two primers together. In contrast, we observed no activity of LigB under these conditions, in agreement with earlier reports that showed a 100-times reduced activity of LigB compared to LigA^39^. In addition, no ligase activity by neither LigA nor LigB is observed on any of the RNA downstream primers, even when the downstream primer contains only a single RNA nucleotide in agreement with early work on *E. coli* ligase activity^40^.

**Fig. 5.**
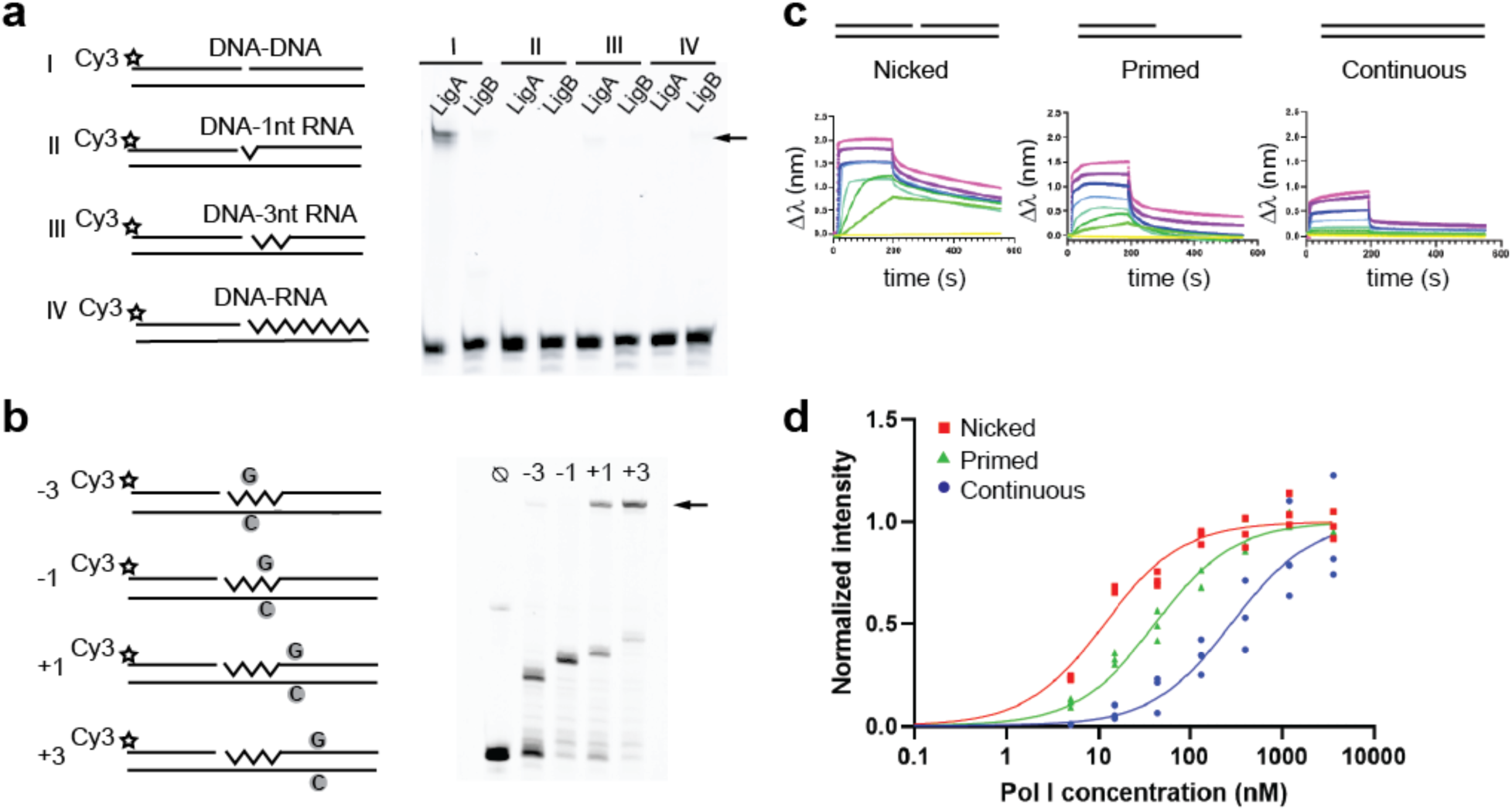
Ligase activity of the *E. coli* LigA and LigB. **a**, Left panel, different DNA-DNA and DNA-RNA substrates used for the ligation assay. Right panel, ligase activity of LigA and LigB on the different DNA/RNA substrates. Arrow marks the full-length ligated product. **b**, Left panel, DNA-RNA substrates with defined stop sites in the RNA and DNA section of the downstream primer. Stop sites are marked with ‘G’ and ‘C’ in a grey circle. Right panel, primer extension and ligation activity of Pol I and LigA on the different DNA/RNA substrates with defined stop sites. Arrow marks the full-length ligated product. **c**, Bio-layer interferometry association and dissociation curves of Pol I on three DNA substrates: a nicked substrate with both an extended and displaced primer, a primed substrate with only an extended primer and a continuous substrate with both template and primer strand of equal length. Different colors indicate increasing protein concentration. **d**, DNA binding curves of Pol I to the three DNA substrates derived from the bio-layer interferometry curves. Data points mark the value at t=180 when association has saturated. Data points are derived from three independent experiments and are normalized to maximum binding value. Derived *K*d values for the different DNA substrates are: nicked 12 ± 5 nM, primed: 80 ± 8 nM, continuous: 800 ± 10 nM.

Thus, the lack of ligase activity on a DNA:RNA junction prevents the incorporation of RNA into the DNA. However, this raises the question: how is Pol I prevented from continuing strand displacement synthesis too far into the DNA section past the RNA primer? We therefore wondered if there may be a role for LigA to evict the polymerase from DNA once the RNA primer has been removed. To determine if this is the case, we designed a series of RNA/DNA substrates that contain a stop position for the polymerase at defined sites in either the RNA or DNA section of the downstream primer. For this, we made use of the high-fidelity DNA synthesis of Pol I that cannot create, or extend from, a mismatched base. By positioning a single C in the template strand and omitting the complementary dGTP from the nucleotide mix during the polymerase assay, we create stop sites that Pol I cannot bypass (Extended Data Fig. 3). The stop positions were chosen at two sites in the RNA primer, at -3 or -1 nucleotides from the RNA/DNA junction, and at two sites in the DNA primer, at +1 or +3 from the RNA/DNA junction (Fig. 5b). All reactions were performed in the presence of LigA, Pol I and the three nucleotides dATP, dCTP, and dTTP. On both substrates where the single C is placed in the RNA section (−3 and -1), we find that the polymerase comes to a halt and cannot proceed. The remaining downstream primer is not ligated to the extended primer, consistent with the observation that LigA cannot ligate DNA to RNA (Fig. 5a). In contrast, when the stop position is located in the DNA section (+1 or +3), we also observe a pause of the polymerase, but the remaining product is now ligated to the extended primer to produce a full-length primer strand. Hence, only when the polymerase reaches the DNA section of the downstream primer, LigA can close the nick.

Next, to determine how the ligation of a nicked substrate impacts the binding of the polymerase, we used bio-layer interferometry to measure the affinity of Pol I for three different DNA substrates: a primed, a nicked, and a continuous DNA substrate (Fig. 5c-d). To prevent end-binding by the polymerase, the free ends of the DNA substrates were blocked by streptavidin. The polymerase binds with high affinity to a nicked substrate (*K*d = 12 ± 5 nM), with a 3.5-fold lower affinity for a primed substrate (*K*d = 42 ± 5 nM), and a 20-fold lower affinity for a continuous stretch of DNA (*K*d = 285 ± 5 nM) (Fig. 5d), implying that as soon as the nick is sealed by LigA, the polymerase will no longer be able to engage with the DNA. Hence, LigA plays a crucial role in discriminating between the RNA and DNA section of the downstream primer and the release of Pol I from the DNA. In combination with the RNA/DNA-junction specific incision of the endonuclease at the end of the RNA primer, it makes for an efficient removal of the RNA primer and prevents the costly continuation of the strand displacement into the DNA section of the downstream Okazaki fragment.

### The β-clamp does not play a role in Okazaki fragment maturation

Finally, we investigated the role of the DNA sliding clamp β during removal of the RNA primer. The β-clamp is essential during DNA replication, where it binds to Pol IIIα and greatly enhances its processivity^10,11^. When Pol IIIα terminates synthesis on a downstream RNA primer and dissociates, the clamp remains bound to the DNA^41–43^ and becomes available for Pol I and/or DNA ligase to bind to. However, contrasting reports exist about the role of the β-clamp during Okazaki fragment maturation. While early reports show a stimulating effect of the β-clamp on the polymerase activity of Pol I^44^, more recent work shows a negative effect^45^. Similarly, early work indicates an interaction between the *E. coli* ligase and β-clamp^44^, while more recently it was reported that the *Mycobacterium tuberculosis* ligase and β-clamp do not interact^46^. Also, in our preliminary analysis, we did not observe an influence of the β-clamp on the activity of Pol I or LigA. This contrast with the eukaryotic system, where the three proteins Pol δ, FEN1, and ligase depend on the eukaryotic clamp PCNA^47–50^. Therefore, to determine if the β-clamp influences the activity of either protein, we designed a DNA substrate that can discriminate between polymerase and ligase activity by using a downstream primer that is longer than the template strand (Fig. 6a). As a result, ligation of the upstream and downstream primer will give a longer product than primer extension of the upstream primer by the polymerase. With increasing amounts of the β-clamp we do not observe a change in the ratio between the extended or the ligated product, indicating that the β-clamp does not affect the activity of the two proteins. Next, we also analyzed the direct interaction between the β-clamp and Pol I or LigA by size exclusion chromatography followed by SDS-PAGE. As judged by the measured SDS-PAGE band intensity of the protein of the elution peak shown in Fig. 6b, the migration of the β-clamp is unaltered by the presence of Pol I or LigA. In contrast, we find a clear shift in the retention volume of the β-clamp in the presence of Pol IIIαε that together firmly bind to the clamp^51,52^. Finally, when mapping the predicted β-binding motifs on Pol I and LigA, we find that they are located in regions of the proteins that are not accessible to the clamp (Extended Data Fig. 4). Taken together, our data indicates that the β-clamp does not play a role in Okazaki fragment maturation in *E. coli*.

**Fig. 6.**
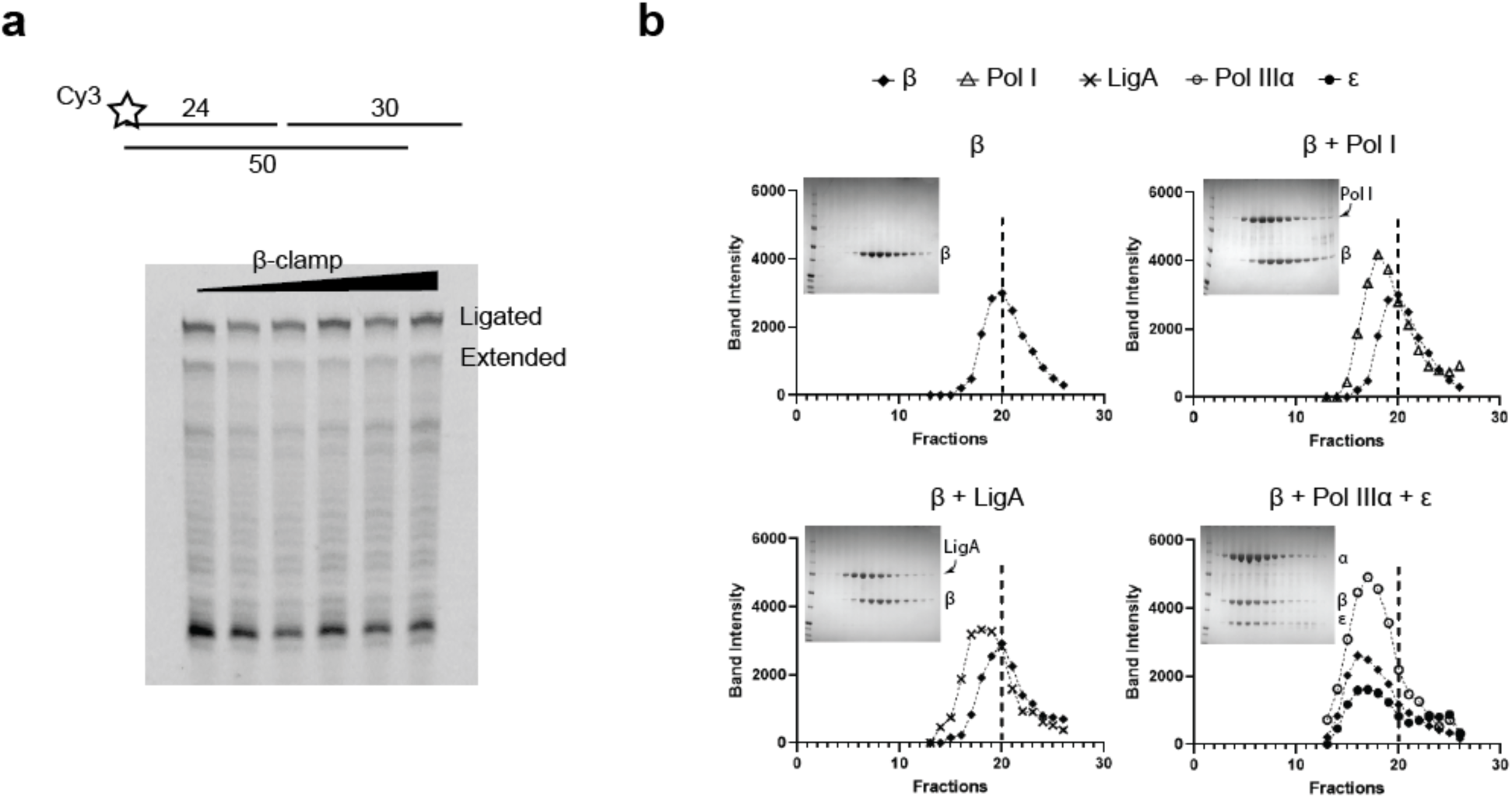
The β-clamp does not interact with Pol I or LigA. **a**, Top: DNA substrate used for primer extension assay. The displaced primer is four nt longer than the template strand so that the ligated product can be separated from the extended product that will be of the same size as the template strand. Bottom: Primer extension assay performed with increasing amounts of β-clamp (0-1.8 μM). **b**, Analytical size exclusion chromatography of β-clamp in the presence of Pol I, LigA and Pol IIIαε. All proteins were injected at 10 μM. Fractions of the chromatography run were analyzed by SDS-page (inserts) and protein band intensities quantified. Graphs show intensities of the individuals protein bands, plotted by fraction number. Dashed vertical line indicates the peak for the isolated β-clamp at fraction 20.

## DISCUSSION

During DNA replication, thousands of RNA primers are incorporated into the lagging strand that subsequently need to be removed to create an intact genome. This is achieved by a series of proteins that together promote the removal of the RNA primer, resynthesis of the DNA, and finally ligation of two adjacent Okazaki fragments. To ensure that this process is performed error free and without spending unnecessary energy, it is essential that the process is well orchestrated. While the activities of the individual proteins have been described, to our knowledge, there has not been a comprehensive analysis of the sequential action of the different proteins. Here, we show that each protein acts as a team member in a molecular relay race, where each protein prepares the DNA/RNA substrate for the next protein to act on (Fig. 7). i) Pol IIIαε synthesizes DNA all the way up to the RNA primer, leaving only a nick. ii) as LigA cannot ligate a DNA-RNA junction, Pol I can enter the DNA and displace the RNA primer while extending the upstream DNA fragment. iii) Pol I endonuclease domain specifically nicks the downstream primer at the RNA/DNA junction to mark the end of the RNA primer. iv) Once all of the RNA primer has been removed, LigA will be able to ligate the nick, preventing further association of the polymerase and leaving an intact DNA fragment. This way, the activities of Pol IIIα, Pol I, endonuclease, and LigA act as a four-point molecular relay race in which each activity is well-coordinated to ensure a fast and efficient removal of the thousands of RNA primers incorporated into the lagging strand during DNA synthesis.

**Fig. 7.**
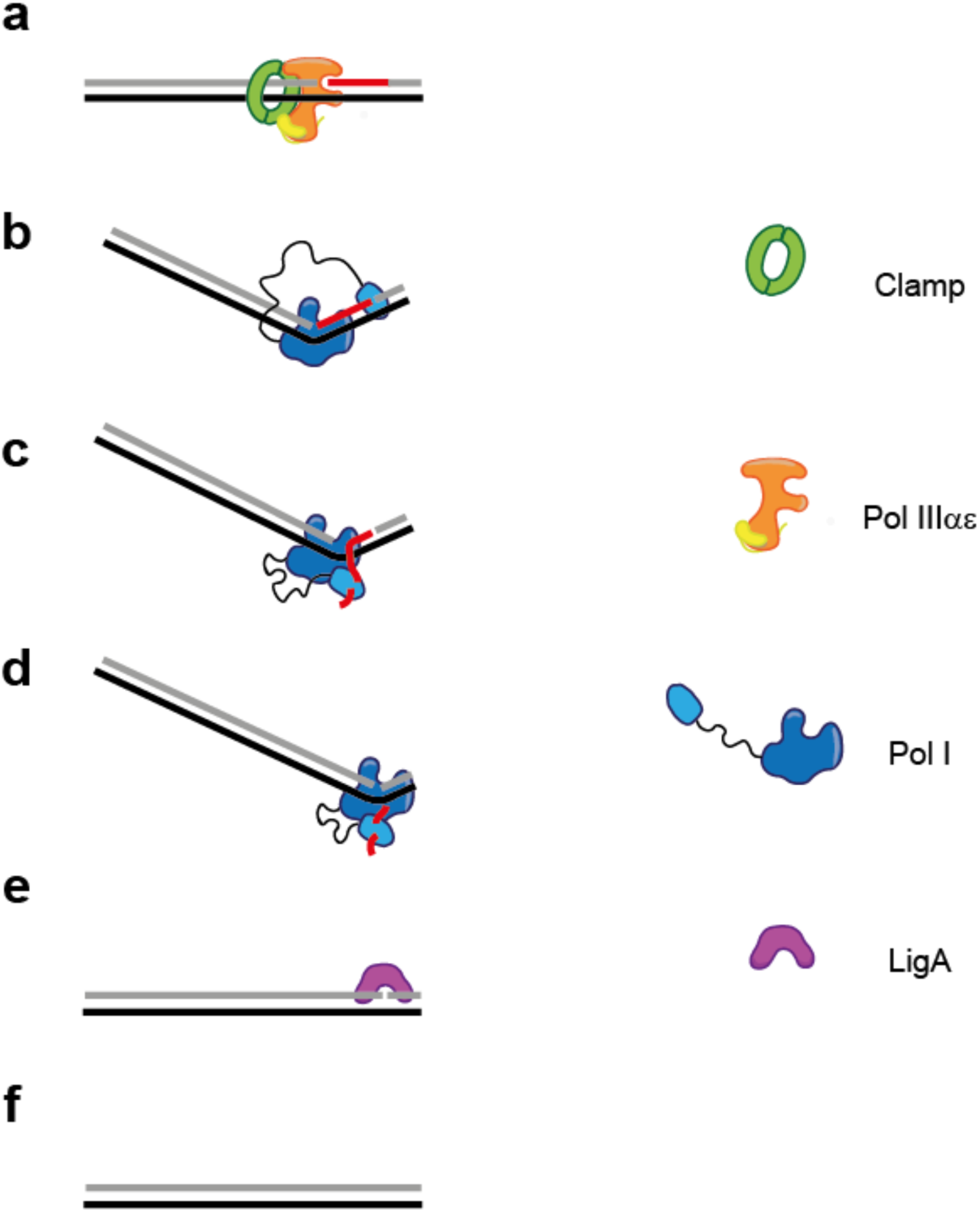
Schematic representation of Okazaki fragment maturation. **a**, The replicative polymerase Pol IIIα, bound to the β-clamp and ε-exonuclease will synthesize up to the last nucleotide before the RNA primer where it will fall off. **b**, Pol I will then bind to the DNA/RNA substrate and kink the DNA to facilitate strand displacement DNA synthesis. In addition, the endonuclease domain will incise the downstream primer at the RNA/DNA junction and mark the end of the RNA primer. **c**, Pol I will then extend the newly synthesized strand, while at the same time displacing the downstream RNA primer. **d**, The displaced strand is cleaved by the endonuclease domain until all of the RNA primer has been removed. **e**, The DNA-DNA junction becomes a substrate for the DNA ligase LigA that will close the gap, **f**, leaving a continuous DNA strand.

Similar to the bacterial system, eukaryotic DNA replication also uses RNA primers that need to be removed from the lagging strand after DNA replication, albeit using a slightly different approach. The lagging strand DNA polymerase δ itself contains modest strand displacement properties^53–55^, while the leading strand polymerase Pol ε does not^53,56^. Yet, unlike Pol I that can remove large stretches of RNA or DNA, Pol δ can only displace 1-2 nucleotides after which it’s 3’-5’ proofreading exonuclease will resect the newly synthesized nucleotides, resulting in ‘idling’ of the polymerase at the nick^55,57^. Therefore, the action of Flap endonuclease 1 (FEN1) is required to remove the displaced flap and allow Pol δ to continue its modest strand displacement synthesis^9,58^. FEN1 distinguishes itself from the bacterial endonuclease in two ways. Unlike the bacterial endonuclease that is part of the same polypeptide, the polymerase FEN1 is an isolated protein that requires the interaction with the DNA sliding clamp PCNA for optimal activity^49^. In addition, while the *E. coli* Pol 1 endonuclease domain can cut flaps of many different sizes (Fig. 3g), FEN1 shows specificity for 1-2 nucleotide flaps^55,59^. Hence, the modest 1-2 nucleotide strand displacement properties of Pol δ fits well with the preference of FEN1 for short flaps. Like the bacterial system, the process is finalized by the sealing of the nick by a DNA ligase LIG1. LIG1, like the *E. coli* LigA, will only ligate DNA to DNA^24^ and therefore prevent incorporation of the RNA primer into the continuous DNA strand. Unlike *E. coli* LigA though, LIG1 relies on the eukaryotic sliding clamp PCNA for its activity^47^. In addition, once LIG1 has sealed the nick, it remains bound to the DNA and PCNA^60^, although the reason for this remains unclear.

Finally, the unique strand displacement DNA synthesis of Pol I is the basis for Loop Mediated Isothermal Amplification (LAMP) that is used for rapid detection of DNA^61^. In combination with a reverse transcriptase (RT-LAMP) the same approach can be used for the amplification of RNA, as used for the detection of SARS-CoV-2 virus^62,63^. Due to the isothermal reaction conditions, LAMP reactions can deliver positive results in under ten minutes^64^, compared to three hours for traditional PCR-based detection methods^65^. The isothermal reaction condition furthermore removes the requirement for specialized equipment, making it possible to employ this detection method in remote places and less developed areas of the world^66^. Our structure of Pol I engaged with both the extended and displaced primer will be of value to further the developments of the enzymes used in the LAMP reaction in order to shorten reaction times, increase sensitivity and accuracy and create a fast, sensitive and reliable assay that can be used all over the world.

## Acknowledgements

We thank the staff of the LUMC EM facility and The Netherlands Center for Electron Nanoscopy (NeCEN) for help with data collection and data processing. This work has been supported by a LUMC Research Fellowship to M.H.L. Access to NeCEN was supported by the Netherlands Electron Microscopy Infrastructure (NEMI), project 184.034.014 of the National Roadmap for Large-Scale Research Infrastructure of the Dutch Research Council (NWO).

## Author contributions

M.H.L. and M.B. conceived the overall experimental design; M.B. prepared samples, performed biochemical assays and analysed data. A.B. collected and processed cryo-EM data; M.B. and M.H.L. wrote the manuscript with contributions from all authors.

## Competing interests

The authors declare no competing interest.

### Reporting Summary

Further information on research design is available in the Nature Research Reporting Summary linked to this article.

## Data availability

Cryo-EM maps and atomic models have been deposited in the Electron Microscopy Database and Protein Data Bank, respectively, under accession codes EMD-nnn, PDB nnn, EMD-ooo and PDB ooo. Source data for graphs shown in Figure 5 are available with the paper online. Other requests should be addressed to Meindert Lamers.

## METHODS

### Chemicals and reagents

All chemicals were purchased from Sigma, unless indicated otherwise. DNA and RNA oligonucleotides were purchased from IDT. Chromatography columns were purchased from Cytiva.

### Mutagenesis

Primers used for generation of expression plasmids and mutagenesis are listed in Supplementary Table 1. The *E. coli* Pol I gene was inserted into a pETNKI-his3C-LIC-kan vector^67^. Pol I Klenow fragment (residues 328-928) that contains only the 3’-5’ exonuclease domain and polymerase domain was cloned into a pETNKI-his3C-LIC-kan vector. The Pol I endonuclease domain (residues 1 to 291) was created by inserting a stop at codon 292 in the vector of full-length Pol I. The endonuclease dead version of full-length Pol I was created by mutagenesis of aspartates 115 and 140 that are analogous to residues 130 and 155 of *M. smegmatis* Pol I that are part of the catalytic site and are required for activity^31^.

**Table 1.** Cryo-EM data collection, refinement, and validation statistics.

| **Table 1. Cryo-EM data collection, refinement, and validation statistics** |  |  |  |
| --- | --- | --- | --- |
| **Data collection and processing** | Pol I<sup>FL</sup> | Pol I<sup>KL</sup> - open | Pol I<sup>KL</sup> - closed |
| Magnification | 105,000 | 105,000 | 105,000 |
| Voltage (kV) | 300 | 300 | 300 |
| Electron exposure e<sup>-</sup>/Å<sup>2</sup> | 54 | 50 | 50 |
| Defocus range (μm) | 0.8 to 2.0 | 0.8 to 2.0 | 0.8 to 2.0 |
| Pixel size (Å) | 0.836 | 0.836 | 0.836 |
| Symmetry imposed | C1 | C1 | C1 |
| Initial particle images (no) | 800892 | 565078 | 565078 |
| Final particle images (no) | 95850 | 164102 | 65904 |
| Map resolution (Å) | 4.3 | 4.0 | 4.1 |
| FSC threshold | 1.43 | 1.43 | 1.43 |
| Map resolution range (Å) | 4.3 to ~ 10 | 4.0 to ~ 10 | 4.1 to ~ 10 |
| **Refinement** |  |  |  |
| Initial model used | 1KLN | 1KLN | 1KLN |
| Model resolution (Å) | 4.3 | 4.0 | 4.1 |
| FSC threshold | 0.143 | 0.143 | 0.143 |
| Map sharpening B factor (Å<sup>2</sup>) | -139 |  |  |
| **Model comparison** |  |  |  |
| Nonhydrogen atoms | 5864 | 5829 | 5891 |
| Protein residues | 603 | 604 | 604 |
| DNA residues | 53 | 51 | 54 |
| **B factors (Å<sup>2</sup>)** |  |  |  |
| Protein | 52-162 | 32-148 | 73-186 |
| **r.m.s deviations** |  |  |  |
| Bond lengths (Å<sup>2</sup>) | 0.004 | 0.009 | 0.007 |
| Bond angles (°) | 1.164 | 1.059 | 0.831 |
| **Validation** |  |  |  |
| MolProbity score | 1.81 | 2.43 | 2.48 |
| Clashscore | 6.56 | 23.81 | 24.93 |
| Poor rotamers (%) | 1.77 | 0 | 0 |
| **Ramachandran plot** |  |  |  |
| Favored (%) | 96.17 | 89.53 | 88.37 |
| Allowed (%) | 3.83 | 10.47 | 11.47 |
| Disallowed (%) | 0 | 0 | 0.17 |

### Protein expression and purification

All proteins were expressed in *E. coli* BL21 (DE3) (Novagen) for two hours at 37 °C, except for LigB that was expressed at 17 °C overnight. All cell pellets were lysed by sonication and clarified by centrifugation at 24,000 x g. Protein purification was performed using buffer A (25 mM Imidazole pH 7.5, 500 mM NaCl, 2mM DTT, 5% Glycerol), buffer B (25mM Tris pH 8.0, 2 mM DTT, 10% Glycerol), buffer C (HEPES pH 7.5, 100 mM NaCl, 2mM DTT), and buffer D (25mM Tris pH 8.5, 2 mM DTT, 10% glycerol), with addition of imidazole or NaCl as indicated. All purified proteins were flash frozen in liquid nitrogen and stored at -80 °C.

Pol I, Pol I^KL^ and Pol I^endo-mut^, Pol IIIα, and β-clamp were purified using a HisTrap column pre-equilibrated in buffer A and eluted using a gradient to 500 mM Imidazole in buffer A. The his-tag was removed by overnight digestion with PreScission Protease (Cytiva), buffer exchanged to buffer A (at 25 mM Imidazole) and followed by a second HisTrap column to remove undigested protein. The flowthrough was injected onto a HiTrap Q column pre-equilibrated in buffer B with 150 mM NaCl and eluted with a gradient to 1 M NaCl in buffer B. Pol IIIα was further purified using a Superdex 200 size exclusion column pre-equilibrated in buffer C.

The endonuclease domain of Pol I was purified using a HisTrap column pre-equilibrated in buffer A and eluted using a gradient to 500 mM Imidazole in buffer A. The eluted protein was diluted 10-fold to 50 mM NaCl in buffer B and injected onto a HiTrap Q column pre-equilibrated in buffer B with 50 mM NaCl and eluted with a gradient to 1 M NaCl in buffer B. The eluted protein was subsequently injected onto a Superdex 200 size exclusion column pre-equilibrated in buffer C.

LigA and LigB were purified using a HisTrap column pre-equilibrated in buffer A and eluted using a gradient to 500 mM Imidazole in buffer A. The eluted protein was diluted 5-fold to 100 mM NaCl in buffer B, injected onto a HiTrap Heparin column pre-equilibrated in buffer B with 100 mM NaCl and eluted with a gradient to 1 M NaCl in buffer B. LigB was subsequently injected onto a Superdex 200 size exclusion column pre-equilibrated in buffer C. LigB elutes in two peaks, which show comparable ligase activity (data not shown). However, the first peak of LigB shows exonuclease activity and was not used for further experiments.

ε is expressed in the insoluble fraction and therefore after cell lysis and centrifugation, the pellet was solubilized in buffer A with 6 M urea and centrifuged at 24,000 x g. The supernatant was injected onto a HisTrap column pre-equilibrated in buffer A with 6 M urea and eluted using a gradient to 500 mM Imidazole in buffer A with 6 M urea. The eluted protein was diluted to 0.5 mg/ml in 25 mM Tris pH 8.5, 3M Urea and 2mM DTT and refolded by overnight dialysis to buffer D. The refolded protein was centrifuged at 24,000 x g and injected onto a HiTrap Q column pre-equilibrated in buffer D with 40 mM NaCl and eluted with a gradient to 1 M NaCl in buffer D.

### Primer extension assays

Primer extension assays were performed using oligonucleotides shown in Supplementary Table 2. All assays were performed in 20 mM HEPES pH 7.5, 2 mM DTT, 5 mM MgCl_2_, 50 mM NaCl and 0.05 mg/ml BSA. For gel analysis, reactions were performed at 22 °C with 100 nM protein and 100 nM of DNA substrate. Primer extensions were carried out for 90 seconds in the presence of 25 μM dNTP’s each. Reactions were stopped in 35 mM EDTA and 65 % formamide and separated on a denaturing 20% acrylamide/bis-acrylamide (19:1) gel with 7.5 M Urea in 1x TBE for 80 minutes at 30 W. The gel was imaged with a Typhoon Imager (GE Healthcare). All experiments were repeated three or more times in independent replicates.

### Cryo-EM sample preparation and imaging

100 μL of purified Pol I^endo-mut^ or Pol I^KL^ at 40 μM was injected onto a 2.4 mL Superdex 200 Increase (3.2/30) column in 100 mM NaCl, 20 mM HEPES pH 7.5 and 2 mM DTT. The peak fraction was collected and incubated for 10 minutes on ice with 20 μM flapped DNA in 20 mM Tris pH 8.5, NaCl 150 mM, 5mM MgCl_2_, 2mM DTT, 0.01% Tween 20 (template: 5’-CGCTCACTGGCCGTCGTTTTACAACGTCGTGACTGGGAAAACCCTGGCGTTACC-3’, upstream primer: 5’-GGTAACGCCAGGGTTTTCCCAGTC -3’, downstream primer: 5’-TTTTTTACGACGTTGTAAAACGACGGCCAGTGAGCG-3’). Three μL of sample at 5 μM was adsorbed onto glow-discharged copper Quantifoil R2/1 holey carbon grids (Quantifoil). Grids were glow discharged 45 seconds at 25 mA using an EMITECH K950 apparatus. Grids were blotted for 1 second at ∼80% humidity at 4°C and flash frozen in liquid ethane. Cryo-EM grids were prepared using a Leica EM GP plunge freezer. The grids were loaded into a Titan Krios (FEI) electron microscope operating at 300 kV with a Gatan K3 detector. The slit width of the energy filter was set to 20 eV. Images were recorded with EPU software (https://www.fei.com/software/epu-automated-single-particles-software-for-life-sciences/) in counting mode. Dose, magnification and effective pixel size are detailed in Table 1.

### Cryo-EM image processing

All image processing was performed using Relion 3.1^68^. The images were drift corrected using Relion’s own (CPU-based) implementation of the UCSF motioncor2 program, and defocus was estimated using CTFFIND4.1^69^. LoG-based auto-picking was performed on a subset of micrographs, and picked particles were 2D classified. Selected classes from the 2D classification were used as references to autopick particles from the full data sets. After three rounds of 2D classification, classes with different orientations were selected for initial model generation in Relion. The initial model was used as reference for 3D classification into different classes. The selected classes from 3D classification were subjected to 3D auto refinement followed by Bayesian polishing. Polished particles were used for 3D classification. Selected particles were subjected to different rounds of CTF refinement plus a final round of Bayesian polishing. Polished particles were used for 3D auto-refine job and the final map was post-processed to correct for modulation transfer function of the detector and sharpened by applying a negative B factor, manually set to -70. A soft mask was applied during post processing to generate FSC curves to yield a map of average resolution of 4.3 Å. The final RELION postprocessed map was used to generate improved-resolution EM maps using the SuperEM method^70^, which aided in model building and refinement.

Model building was performed using Coot^71^, REFMAC5^72^, the CCPEM-suite^73^. Details on model refinement and validation are in Table 1. In brief, model building started by rigid-body fitting of the known *E. coli* Pol1 Klenow crystal structure^28^ (PDB 1KLN) into experimental density map using Coot. The DNA molecule was generated in Coot and rigid body fitted into experimental density map. Next, we carried out one round of refinement in Refmac5 using jelly-body restraints, and the model was further adjusted in Coot. A final refinement round and validation of the model and data were carried out using Refmac5 with proSmart^74^ restraints and MolProbity^75^ within the CCPEM suite.

### Analytical size exclusion chromatography

Samples of the different complexes were prepared at 10 μM and 100 μl were injected onto a 2.4 mL Superdex 200 Increase gel filtration column (GE Healthcare) pre-equilibrated in 100 mM NaCl, 20 mM HEPES pH 7.5 and 2 mM DTT. For all, 70 μl fractions were collected and analysed by SDS-PAGE using 4-12% NuPAGE Bis-Tris gels (Invitrogen). The gels were run in MOPS buffer at 200 V for 45 minutes and stained with InstantBlue Coomassie protein stain (Abcam). Protein bands intensities were quantified using ImageJ^76^ and then plotted against fraction numbers using GraphPad Prism version 9.0.1.

### Sequence analysis of β-binding motif

The canonical β-binding motif Q-x(2)-[LM]-x-F and Q-x(2)-[LM]F^77^ was used to search the sequences of *E. coli* Pol I and LigA using ScanProsite^78^. Pol I contains one potential β-binding motif (residues 587-591), while LigA contains two potential β-binding motifs (residues 297-302 and 318-323).

### Bio-layer Interferometry

The Octet RED96 instrument (ForteBio) was used for bio-layer interferometry data acquisition. Three different DNA substrates were prepared for the assays (sequences are shown in Supplementary Table 3). A 5’ biotinylated primer and monovalent streptavidin^79^ were mixed in a 1:1 ratio at a concentration of 20 μM and purified via analytical gel filtration using a Superdex 200 increase (3.2/30) column equilibrated in 10 mM Tris pH 8.0, 50 mM NaCl, and 1 mM EDTA EDTA 1mM. Fractions containing biotinylated primer bound to monovalent streptavidin were subsequently annealed to a 5’ biotinylated DNA template before binding to streptavidin biosensors (Sartorius). 200 nM of DNA substrates were added in the loading step, until the threshold value of 0.4 nm was reached. All the experiments were performed in 10 mM Tris pH 7.5, 150 mM KCl, 5 mM MgCl_2_, 0.05% Tween 20%, 2 mM DTT, 0.1% BSA and 1mg/ml Dextran. The association step was performed by a titration of protein concentration starting from 0 to 3.6 μM of Pol I. The dissociation step was performed in buffer alone. Kinetic analysis was performed using the Octet Data Analysis software package version 7.1 and GraphPad Prism version 9.0.1.

