## Supplementary Material for "A four-point molecular handover during Okazaki maturation"

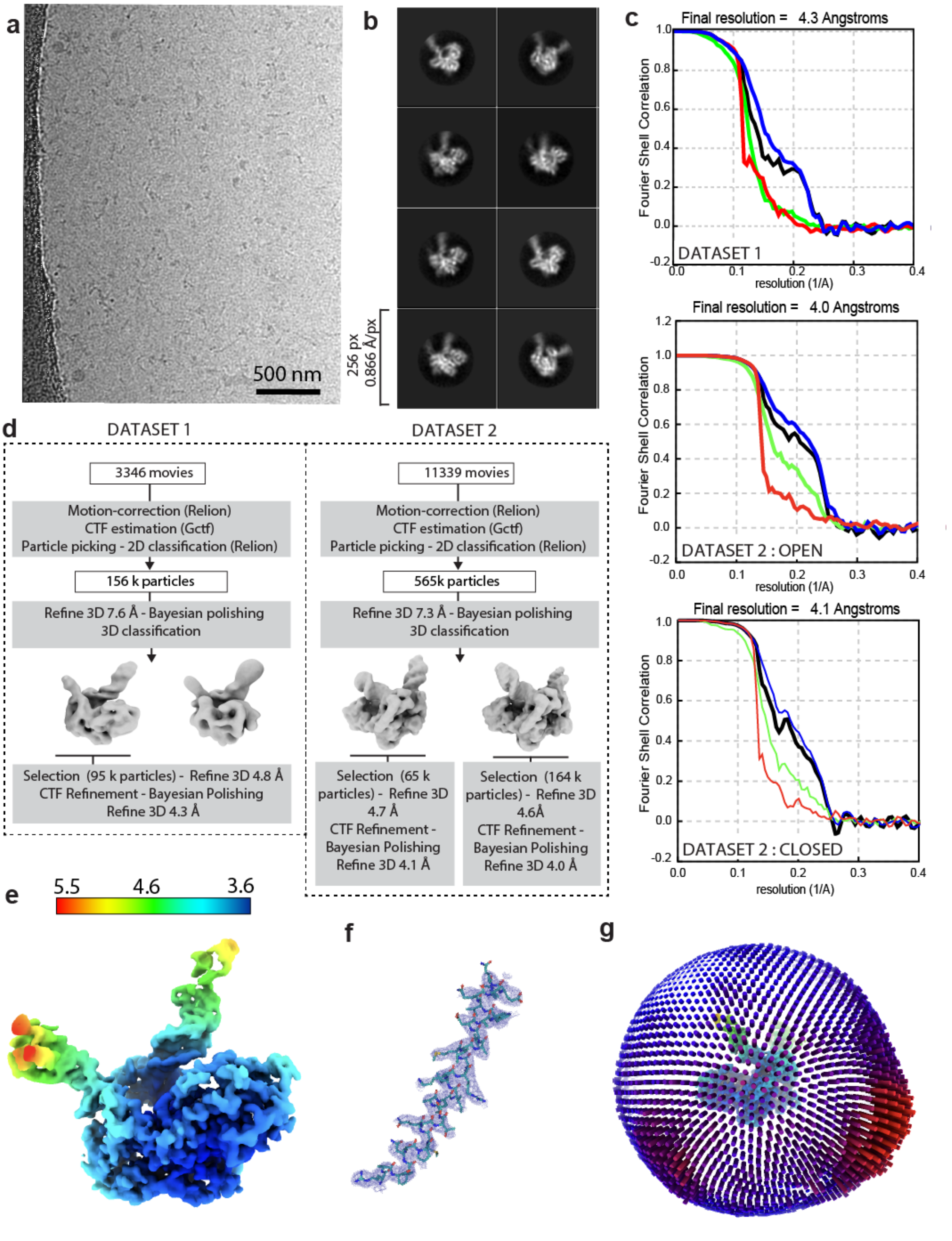

**Extended Data Fig. 1 | Cryo-EM data collection and data processing details**. **a**, Representative micrograph. **b**, 2D class averages from full dataset. **c**, Fourier Shell Correlation between half-maps from final refinement. Green line: unmasked. Blu line: masked. Red line: phase randomized. Black line: corrected. **d**, Schematic representation of main data processing procedures. See methods section for more details. **e**, Final map obtained applying SuperEM code to Relion post-processed map from dataset 1 and colored by local resolution. **f**, Detail of model fitted to the final cryo-EM map from dataset 1. **g**, Orientational distribution of final set of refined particles from dataset 1.

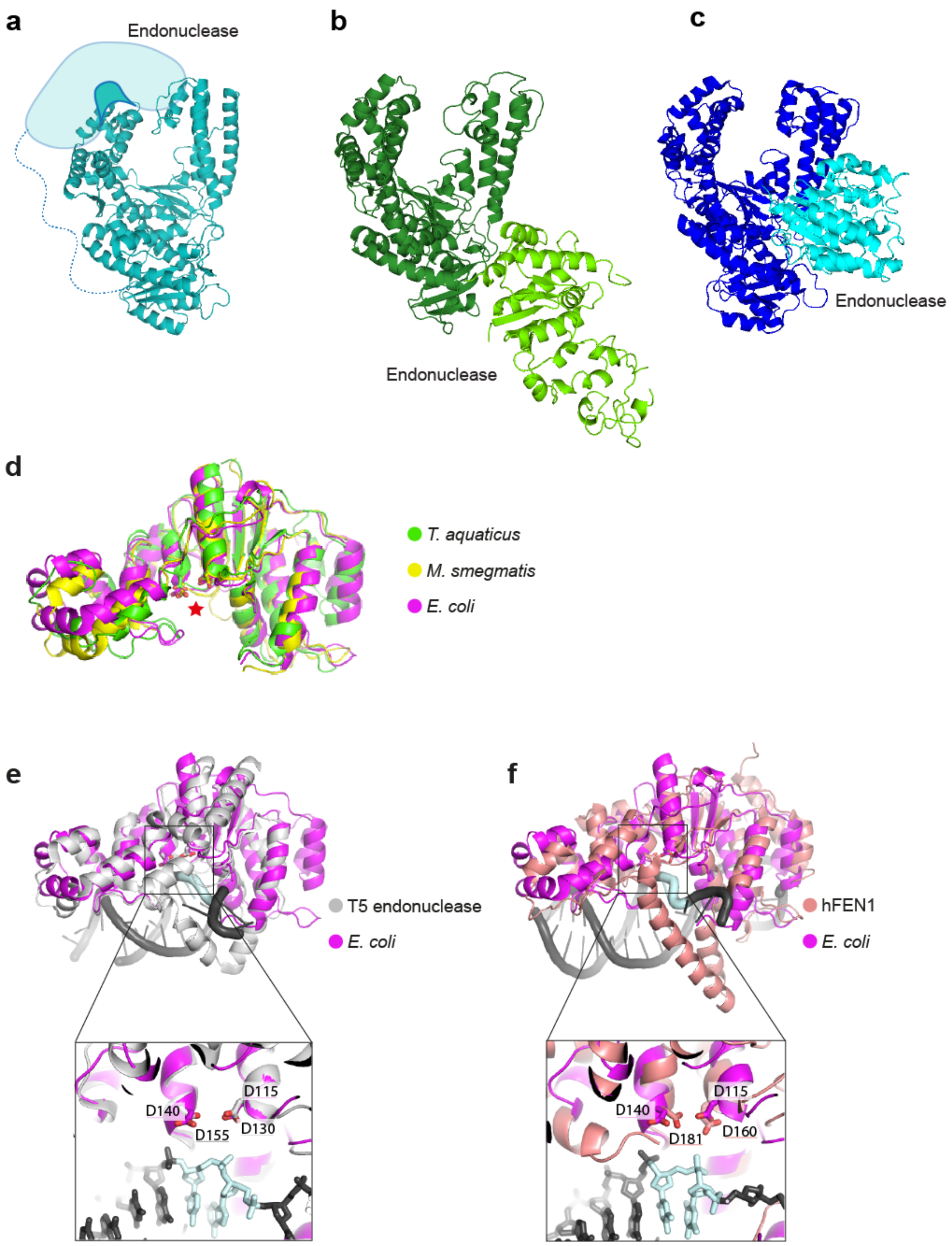

**Extended Data Fig. 2 | Positional modelling of the endonuclease domain in Pol I.** **a**, *E. coli* Pol I with predicted position of the endonuclease domain on top of the fingers domain, based on the position of the single stranded flap (see also main Fig. 3a). **b**, *Thermus aquaticus* Pol I with its endonuclease domain adjacent to the 3'-5'

exonuclease domain[33]. **c**, *Mycobacterium smegmatis* Pol I with its endonuclease domain adjacent to the thumb domain[31]. **d**, Superposition of three Pol I endonuclease domains from *T. aquaticus*, *M. smegmatis*, and *E. coli*, (modelled using AlphaFold[34]). Red star marks the endonuclease active site. **e**, Superposition of *E. coli* Pol I endonuclease domain and T5 endonuclease bound to its substrate DNA[35]. The square marks the enlarged area shown below. Site of incision is marked by the two nucleotides in light blue. Two catalytic aspartates are shown in sticks. **f**, Similar comparison with human FEN1[36]. Structures of T5 endonuclease and FEN1 were used to model the position of Pol I endonuclease in main Fig. 3b.

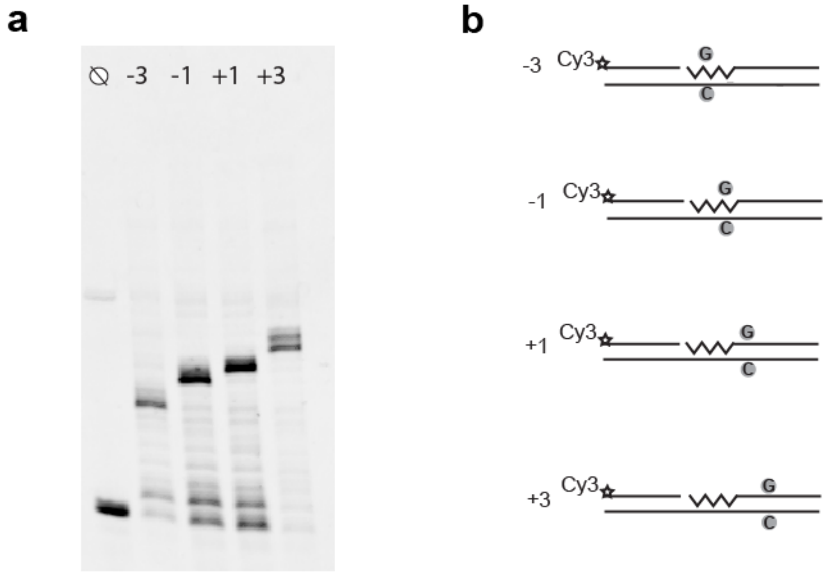

**Extended Data Fig. 3 | Pol I does not progress past a C in the template strand in absence of dGTP. a**, Pol I activity in absence of LigA using the templates shown in panel b, using only the three nucleotides dATP, dCTP, and dTTP. Experimental conditions are the same as in Fig. 5b. **b**, Substrates used for experiment in panel a.

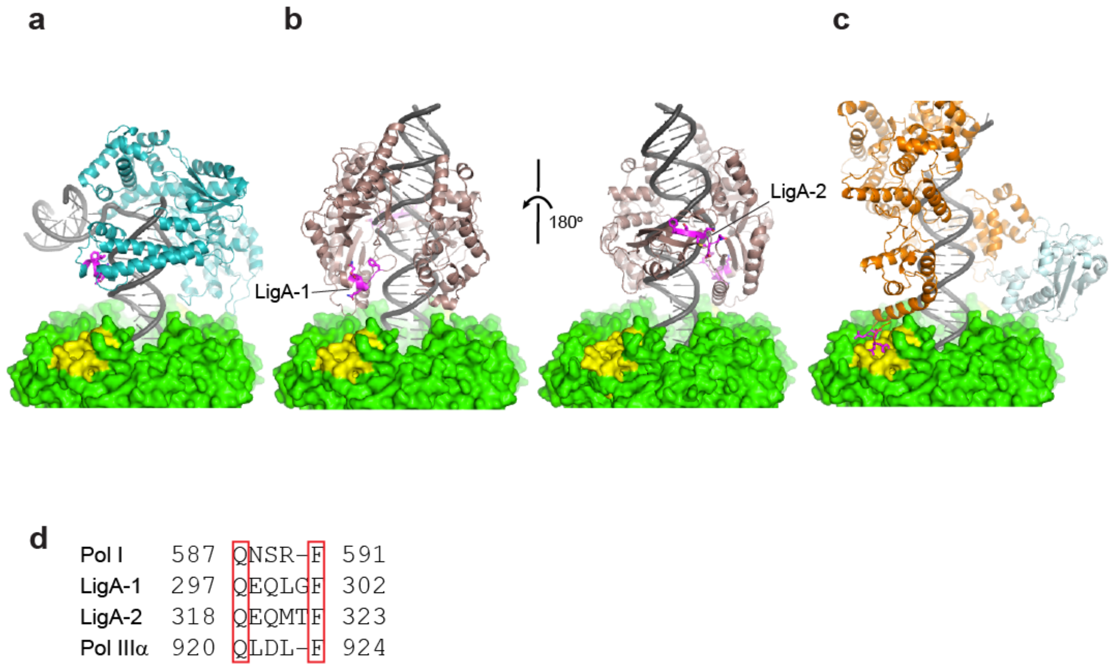

**Extended Data Fig. 4 | The location of β-binding motifs of Pol I, LigA and Pol IIIα.** **a**, Model of Pol I and β-clamp. The β-clamp is shown in green surface, with binding pocket highlighted in yellow. The predicted β-binding motif of Pol I is shown in magenta sticks and is located in a helix of the thumb domain that interacts with the minor groove of the DNA. **b**, Model of LigA[79] and β-clamp shown in two views rotated by 180°. The two predicted β-binding motifs are shown in magenta sticks. Motif LigA-1 is located in a helix that precedes a loop that interacts with the DNA. Motif LigA-2 is located on a strand that is part of the OB-domain that also interacts with the DNA. **c**, Cryo-EM structure of Pol IIIα bound to the β-clamp, exonuclease ε, and DNA[52]. The Pol IIIα β-binding motif is shown in magenta sticks and located on a loop at the end of the fingers domain and interacts with the binding pocket of the β-clamp. **d**, Alignment of β-binding motifs from Pol I, LigA and Pol IIIα that are highlighted in magenta in panels a-c.

| Primer name | Length | 5'-3' sequence |
|---|---|---|
| ASP115ALA Fw | 26 | GCGTAGAAGCGGCCGACGTTATCGGT |
| ASP115ALA Rv | 26 | ACCGATAACGTCGGCCGCTTCTACGC |
| ASP140ALA Fw | 28 | GCACTGGCGATAAAGCTATGGCGCAGCT |
| ASP140ALA Rv | 28 | AGCTGCGCCATAGCTTTATCGCCAGTGC |
| Third Stop Codon Pol1 Fw | 24 | TGCTGATGTCTAAGCGGGCAAATG |
| Third Stop Codon Pol1 Rv | 24 | CATTTGCCCGCTTAGACATCAGCA |

**Supplementary Table 1** Primers used for cloning and mutagenesis

| Figure n° | Panel | Sequences |
|---|---|---|
| 1 | A | 5′ Cy3-GGTAACGCCAGGGTTTTCCCAGTC 3′<br>3′ CCATTGCGGTCCCAAAAGGGTCAGTGAAGTGCTTAGATCTGCTGCAACATTTTGCTGCCGGTCACTCGC 5′ |
| | B | 5′ Cy3-GGTAACGCCAGGGTTTTCCCAGTC ACGACGTTGTAAAACGACGGCCAGTGAGCG 3′<br>3′ CCATTGCGGTCCCAAAAGGGTCAGTGAAGTGCTTAGATCTGCTGCAACATTTTGCTGCCGGTCACTCGC 5′ |
| | C | 5′ Cy3-GGTAACGCCAGGGTTTTCCCAGTC *ACGACGTTGTAAAACGACGGCCAGTGAGCG 3′<br>3′ CCATTGCGGTCCCAAAAGGGTCAGTGAAGTGCTTAGATCTGCTGCAACATTTTGCTGCCGGTCACTCGC 5′ |
| 3 | D | 5′Cy3-GGTAACGCCAGGGTTTTCCCAGTC 3′<br>3′CCATTGCGGTCCCAAAAGGGTCAGTGCTGCAACATTTTGCTGCCGGTCACTCGC 5′ |
| | E | 5′Cy3-GGTAACGCCAGGGTTTTCCCAGTCACGACGTTGTAAAACGACGGCCAGTGAGCG 3′<br>3′CCATTGCGGTCCCAAAAGGGTCAGTGCTGCAACATTTTGCTGCCGGTCACTCGC 5′ |
| | F | 5′GGTAACGCCAGGGTTTTCCCAGTCACGACGTTGTAAAACGACGGCCAGTGAGCG-Cy5 3′<br>3′CCATTGCGGTCCCAAAAGGGTCAGTGCTGCAACATTTTGCTGCCGGTCACTCGC 5′<br><br>A<br>5′GGTAACGCCAGGGTTTTCCCAGTCGCGACGTTGTAAAACGACGGCCAGTGAGCG-Cy5 3′<br>3′ CCATTGCGGTCCCAAAAGGGTCAGCGCTGCAACATTTTGCTGCCGGTCACTCGC 5′<br><br>AC<br>5′ GGTAACGCCAGGGTTTTCCCAGTCGTGACGTTGTAAAACGACGGCCAGTGAGCG-Cy5 3′<br>3′ CCATTGCGGTCCCAAAAGGGTCAGCACTGCAACATTTTGCTGCCGGTCACTCGC 5′<br><br>ACG<br>5′ GGTAACGCCAGGGTTTTCCCAGTCGTAACGTTGTAAAACGACGGCCAGTGAGCG-Cy5 3′<br>3′ CCATTGCGGTCCCAAAAGGGTCAGCATTGCAACATTTTGCTGCCGGTCACTCGC 5′<br><br>ACGA<br>5′ GGTAACGCCAGGGTTTTCCCAGTCGTACCGTTGTAAAACGACGGCCAGTGAGCG-Cy5 3′<br>3′ CCATTGCGGTCCCAAAAGGGTCAGCATGGCAACATTTTGCTGCCGGTCACTCGC 5′<br><br>ACGAC<br>5′ GGTAACGCCAGGGTTTTCCCAGTCGTACGGTTGTAAAACGACGGCCAGTGAGCG-Cy5 3′<br>3′ CCATTGCGGTCCCAAAAGGGTCAGCATGCCAACATTTTGCTGCCGGTCACTCGC 5′ |
| 4 | A | 5′CTTAACTCCAACCTTTTCCCAATCACTTCACGAACCACATTCTAAAACCACTTCCATTCACCT-Cy5 3′<br>3′GAATTGAGGTTGGAAAAGGGTTAGTGAAGTGCTTGGTGTAAGATTTTGGTGAAGGTAAGTGGA 5′<br><br>5′CTTAACTCCAACCTTTTCCCAATC**ACUUCACGAA**CCACATTCTAAAACCACTTCCATTCACCT-Cy5 3′<br>3′GAATTGAGGTTGGAAAAGGGTTAGTGAAGTGCTTGGTGTAAGATTTTGGTGAAGGTAAGTGGA 5′ |
| | B | 5′CTTAACTCCAACCTTTTCCCAATC**ACUUCACGAA**CCACATTCTAAAACCACTTCCATTCACCT-Cy5 3′<br>3′GAATTGAGGTTGGAAAAGGGTTAGTGAAGTGCTTGGTGTAAGATTTTGGTGAAGGTAAGTGGA 5′ |
| | C | 5′CTTAACTCCAACCTTTTCCCAATC**ACUUCACUAA**GCACATTCTAAAACCACTTCCATTCACCT-Cy5 3′<br>3′GAATTGAGGTTGGAAAAGGGTTAGTGAAGTGATTGGCGTAAGATTTTGGTGAAGGTAAGTGGA 5′<br><br>5′CTTAACTCCAACCTTTTCCCAA **ACUUCACUAA**GCACATTCTAAAACCACTTCCATTCACCT-Cy5 3′<br>3′GAATTGAGGTTGGAAAAGGGTTAGTGAAGTGATTGGCGTAAGATTTTGGTGAAGGTAAGTGGA 5′<br><br>5′CTTAACTCCAACCTTTTCCC **ACUUCACUAA**GCACATTCTAAAACCACTTCCATTCACCT-Cy5 3′<br>3′GAATTGAGGTTGGAAAAGGGTTAGTGAAGTGATTGGCGTAAGATTTTGGTGAAGGTAAGTGGA 5′<br><br>5′CTTAACTCCAACCTTTT **ACUUCACUAA**GCACATTCTAAAACCACTTCCATTCACCT-Cy5 3′<br>3′GAATTGAGGTTGGAAAAGGGTTAGTGAAGTGATTGGCGTAAGATTTTGGTGAAGGTAAGTGGA 5′ |

| | | |
|---|---|---|
| 5 | A | 5’Cy3-GGTAACGCCAGGGTTTTCCCAGTCACGACGTTGTAAAACGACGGCCAGTGAGCG 3’<br>3’CCATTGCGGTCCCAAAAGGGTCAGTGCTGCAACATTTTGCTGCCGGTCACTCGC 5’ |
| | | 5’Cy3-GGTAACGCCAGGGTTTTCCCAGTC**A**CGACGTTGTAAAACGACGGCCAGTGAGCG 3’<br>3’CCATTGCGGTCCCAAAAGGGTCAGTGCTGCAACATTTTGCTGCCGGTCACTCGC 5’ |
| | | 5’Cy3-GGTAACGCCAGGGTTTTCCCAGTC**ACG**ACGTTGTAAAACGACGGCCAGTGAGCG 3’<br>3’CCATTGCGGTCCCAAAAGGGTCAGTGCTGCAACATTTTGCTGCCGGTCACTCGC 5’ |
| | | 5’Cy3-GGTAACGCCAGGGTTTTCCCAGTC**ACGACGUUGUAAAACGACGGCCAGUGAGCG** 3’<br>3’CCATTGCGGTCCCAAAAGGGTCAGTGCTGCAACATTTTGCTGCCGGTCACTCGC 5’ |
| | B | 5’Cy3-CTTAACTCCAACCTTTTCCCAATC**ACUUCACGAA**CCACATTCTAAAACCACTTCCATTCACCT 3’<br>3’GAATTGAGGTTGGAAAAGGGTTAGTGAAGTGCTTGGTGTAAGATTTTGGTGAAGGTAAGTGGA 5’ |
| | | 5’Cy3-CTTAACTCCAACCTTTTCCCAATC**ACUUCACUAG**CCACATTCTAAAACCACTTCCATTCACCT 3’<br>3’ GAATTGAGGTTGGAAAAGGGTTAGTGAAGTGATCGGTGTAAGATTTTGGTGAAGGTAAGTGGA 5’ |
| | | 5’Cy3-CTTAACTCCAACCTTTTCCCAATC**ACUUCACUAA**GCACATTCTAAAACCACTTCCATTCACCT 3’<br>3’ GAATTGAGGTTGGAAAAGGGTTAGTGAAGTGATTCGTGTAAGATTTTGGTGAAGGTAAGTGGA 5’ |
| | | 5’Cy3-CTTAACTCCAACCTTTTCCCAATC**ACUUCACUAA**GCACATTCTAAAACCACTTCCATTCACCT 3’<br>3’ GAATTGAGGTTGGAAAAGGGTTAGTGAAGTGATTCGTGTAAGATTTTGGTGAAGGTAAGTGGA 5’ |
| | | 5’Cy3-CTTAACTCCAACCTTTTCCCAATC**ACUUCACUAA**CCGCATTCTAAAACCACTTCCATTCACCT 3’<br>3’ GAATTGAGGTTGGAAAAGGGTTAGTGAAGTGATTGGCGTAAGATTTTGGTGAAGGTAAGTGGA 5’ |
| 6 | A | 5’Cy3-GGTAACGCCAGGGTTTTCCCAGTCACGACGTTGTAAAACGACGGCCAGTGAGCG 3’<br>3’CCATTGCGGTCCCAAAAGGGTCAGTGCTGCAACATTTTGCTGCCGGTCAC 5’ |

**Supplementary Table 2** Primers used for primer extension assays. RNA highlighted in bold. Underlined letters indicate the position of the nick. Phosphorylation of substrate at 5’ end indicated with an asterisk.

| **Substrate name** | **Sequences** |
|---|---|
| Continuous | 5’GGTAACGCCAGGGTTTTCCCAGTCACGACGTTGTAAAACGACGGCCAGTGAGCG-B 3’<br>3’B-CCATTGCGGTCCCAAAAGGGTCAGTGCTGCAACATTTTGCTGCCGGTCACTCGC 5’ |
| Primed | 5’ACGACGTTGTAAAACGACGGCCAGTGAGCG-B 3’<br>3’B-CCATTGCGGTCCCAAAAGGGTCAGTGCTGCAACATTTTGCTGCCGGTCACTCGC 5’ |
| Nicked | 5’GGTAACGCCAGGGTTTTCCCAGTCACGACGTTGTAAAACGACGGCCAGTGAGCG-B 3’<br>3’B-CCATTGCGGTCCCAAAAGGGTCAGTGCTGCAACATTTTGCTGCCGGTCACTCGC 5’ |

**Supplementary Table 3** Primers used for biolayer interferometry assays. Underlined letters indicate the position of the nick. All primers contain a biotin at 3’ end.
